# Compensatory mechanisms affect sensorimotor integration during ongoing vocal-motor acts in marmoset monkeys

**DOI:** 10.1101/696989

**Authors:** Thomas Pomberger, Julia Löschner, Steffen R. Hage

## Abstract

In vertebrates, any transmission of vocal signals faces the challenge of acoustic interferences such as heavy rain, wind, animal, or urban sounds. Consequently, several mechanisms and strategies have evolved to optimize the signal-to-noise ratio. Examples to increase detectability are the Lombard effect, an involuntary rise in call amplitude in response to masking ambient noise, which is often associated with several other vocal changes such as call frequency and duration, as well as the animals’ capability of limiting calling to periods where noise perturbation is absent. Previous studies revealed rapid vocal flexibility and various audio-vocal integration mechanisms in marmoset monkeys. Using acoustic perturbation triggered by vocal behavior, we investigated whether marmoset monkeys are capable of exhibiting changes in call structure when perturbing noise starts after call onset or whether such effects only occur if noise perturbation starts prior to call onset. We show that marmoset monkeys are capable of rapidly modulating call amplitude and frequency in response to such perturbing noise bursts. Vocalizations swiftly increased call frequency after noise onset indicating a rapid effect of perturbing noise on vocal motor pattern production. Call amplitudes were also affected. Interestingly, however, the marmosets did not exhibit the Lombard effect as previously reported but decreased their call intensity in response to perturbing noise. Our findings indicate that marmosets possess a general avoidance strategy to call in the presences of ambient noise and suggest that these animals are capable of counteracting a previously thought involuntary audio-vocal mechanism, the Lombard effect, presumably via cognitive control processes.

## Introduction

Communication between individuals is a crucial aspect for evolutionary success and appears in various forms in nature ranging from olfactory [1,2] to visual [3] to vocal signals [4]. For proper communication, the transmission of a signal sent out by a sender has to be detected and decoded by one or more receivers [5]. Therefore, the sender has to be able to modulate the signal in response to potential masking ambient noise to ensure proper signal transmission. For vocal communication in vertebrates, several mechanisms have evolved to compensate for masking acoustic interferences, such as heavy rain, wind, animal, or urban sounds, leading to changes in temporal and spectral features of the vocal signals [6]. Such vocal modifications can happen involuntarily as well as under volitional control.

One of the most efficient mechanisms to increase signal-to-noise ratio in call production is the so-called Lombard effect, i.e., the involuntary increase in call amplitude in response to masking ambient noise [7]. It is often accompanied by a shift in call frequency[8,9] as well as a change in call duration [10,11] and has been shown in many vertebrate species from fish to frogs to birds to mammals including humans [12,13], suggesting that the Lombard effect is an evolutionary old behavior that may have emerged about 450 million years ago. Another successful strategy to increase detectability in a noisy environment is the restraint of call emission to timeslots where noise perturbation is low or absent [10,14,15]. This approach renders the modification of call parameters unnecessary and avoids the increased physiological cost of call emission at high intensities that might still be insufficiently increasing signal-to-noise ratio.

The common marmoset, a small, highly vocal New World monkey indigenous to the dense rainforests of Brazil, has been shown to exhibit vocal flexibility, such as increasing call intensity [16,17] or increasing the duration of specific calls [17], as well as the attempt to call in silent gaps [14], in the presence of perturbing ambient noise. These findings suggest that while these animals generally seem to prefer avoiding calling in a noisy environment, they do exhibit the involuntary audio-vocal effects discussed above when doing so. This idea is supported by a recent study showing that marmoset tend to produce single calls instead of call sequences in response to perturbing noise stimuli [18]. Interestingly, marmoset monkeys are also capable of interrupting ongoing vocalizations rapidly after noise perturbation onset [18], overturning decades-old concepts regarding vocal pattern generation [19–21], indicating that vocalizations do not consist of one discrete call pattern but are built of many sequentially uttered units that might be modulated and initiated independently of each other. However, it is yet unclear whether audio-vocal mechanisms, such as the Lombard effect and its accompanied changes in call frequency, can be rapidly elicited in cases where the perturbing noise starts after call onset or whether such effects only occur if noise perturbation starts prior to call onset.

In the present study we use acoustic perturbation triggered by the vocal behavior itself to test in a controlled experimental design whether marmosets are capable of rapidly modulating distinct vocal parameters such as call frequency and amplitude in ongoing vocalizations. Performing quantitative measures of resulting adjustments, we show that marmoset monkeys are able to specifically and rapidly modulate call frequency and amplitude as a response to white noise stimuli in ongoing vocal utterances. Hereby, our data indicate that marmosets exhibit a decrease in call amplitude as a result of such noise perturbation, suggesting a mechanism counteracting the rise in amplitude caused by the Lombard effect.

## Results

We measured vocal behavior in marmoset monkeys (*Callithrix jacchus*, n = 4), a highly vocal New World monkey species, while separated in a soundproofed chamber, with and without acoustic perturbation (**Fig. 1A and B**). In this setting, marmoset monkeys predominantly produced phee calls (monkey H: 92.0%, S: 99.1%, F: 96.8%, W: 95.6%), long-distance contact calls, composed of one (so-called single phees), two (double phees), or more phee syllables, to interact with conspecifics [22] (**Fig. 1C**). Other call types such as trill-phees, twitters, trills, tsik-ekks [22,23], and segmented phees [24] were rarely uttered (all other call types were well below 2.5% in all monkeys except trill-phees in monkey H [4.6%]).

**Figure 1:**
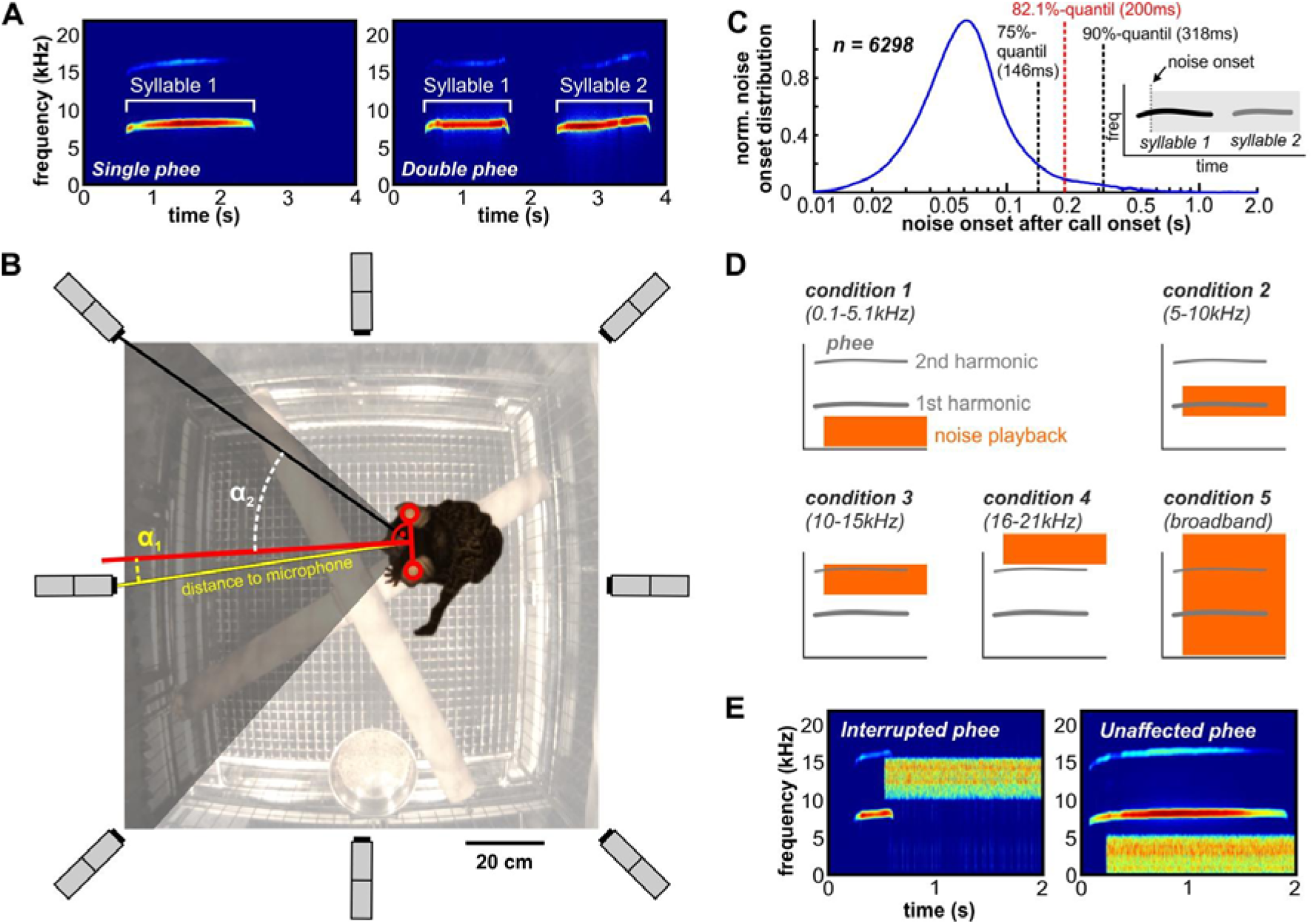
Experimental setup and design. (**A**) Exemplary spectrograms of single and double phee calls. (**B**) The vocal behavior of monkeys was recorded in a soundproof chamber. The behavior was continuously monitored and recorded. The red line shows the monkey’s head position in relation to the two closest microphones (yellow and black line). The acoustic signal recorded with the microphone closest to being directly in front of the monkey’s head (i.e., the smallest angle between the monkey’s perpendicular and the microphone) was used for amplitude calculation. (**C**) Relative vocal detection distribution over time (s). (**D**) Noise condition overview with masking properties. (**E**) Exemplary spectrograms of an interrupted single phee (10–15 kHz noise condition) and unaffected phee (0.1–5 kHz noise condition).

We perturbed 2/3 of calls with noise playback after vocal onset to ensure perturbation starting after call initiation (**Fig. 1B**). To investigate whether perturbation of different frequency bands within the hearing range of the monkeys has different effects on their vocal behavior, we played back five different noise band conditions (broadband noise and bandpass filtered noise bands below [0.1–5 kHz], around [5–10 kHz], or above the fundamental frequency of phee calls [noise bands of 10–15 kHz and 16–21 kHz] at four different amplitudes [50 dB, 60 dB, 70 dB, 80 dB] each). All noise conditions were played back pseudo-randomly in blocks of 30 uttered vocalizations, resulting in 20 calls being perturbed by noise after call onset and 10 calls not being perturbed by noise (control). In total, our monkeys produced 6,298 phees (monkey F = 1544 phees, H = 1471, S = 1631, W = 1652). Monkeys uttered mostly single and double phees (multi-syllabic phees with more than two syllables were rare or absent: monkey F = 6.5%, H = 0.4%, S = 1.3%, absent in W), with double phee rates between 8.4% and 55.5% (mean: 29.5% ± 9.8%, n = 4 monkeys) in the control condition.

We first investigated if and how marmosets changed the fundamental frequency of their ongoing phee syllables when perturbed by different noise conditions. We found an increase in first syllable frequencies (F(3,4904)=6.42, p=2.0e-04 for amplitude, F(4,4904)=20.68, p<0.0001 for frequency, n=3180). Those frequency shifts were significant in the 0.1–5.1 kHz at 80 dB noise condition (38.5±13.8 Hz, p=1.02e-02, n=168), in the two loudest conditions of the 5–10 kHz noise band (70 dB: 56.9±14.5 Hz, p=3.20e-03, n=165; 80 dB: 76.7±16.4 Hz, p=4.12e-08, n=134), and in all four amplitude conditions of broadband noise (50 dB: 39.2±13.7 Hz, p=5.03e-10, n=159; 60 dB: 68.6±13.4 Hz, p=1.21e-08, n=143; 70 dB: 104.1±12.5 Hz, p=3.99e-13, n=135; 80 dB: 101.7±16.1 Hz, p=1.66e-20, n=118; control: n=1733; **Fig. 2A**). The largest frequency shift could be observed for 70 dB broadband noise, while in the next higher intensity condition (80 dB), there was no further increase in frequency (p=1, n=253), indicating that marmosets are only capable of altering their fundamental frequency within a certain range. Frequency shifts were not observed in calls that were produced during 10–15 kHz and 16–21 kHz noise band perturbations (p=1, n=669 for the 10–15 kHz noise band, n=652 for the 16–21 kHz noise band, **Fig. 2A**). Second phee syllables showed no significant shift in fundamental frequencies when perturbed by noise (F(3,1343)=1.56, p=0.20 for amplitude, F(4,1343)=1.24, p=0.29 for frequency, n=761, **Fig. 2B**).

**Figure 2:**
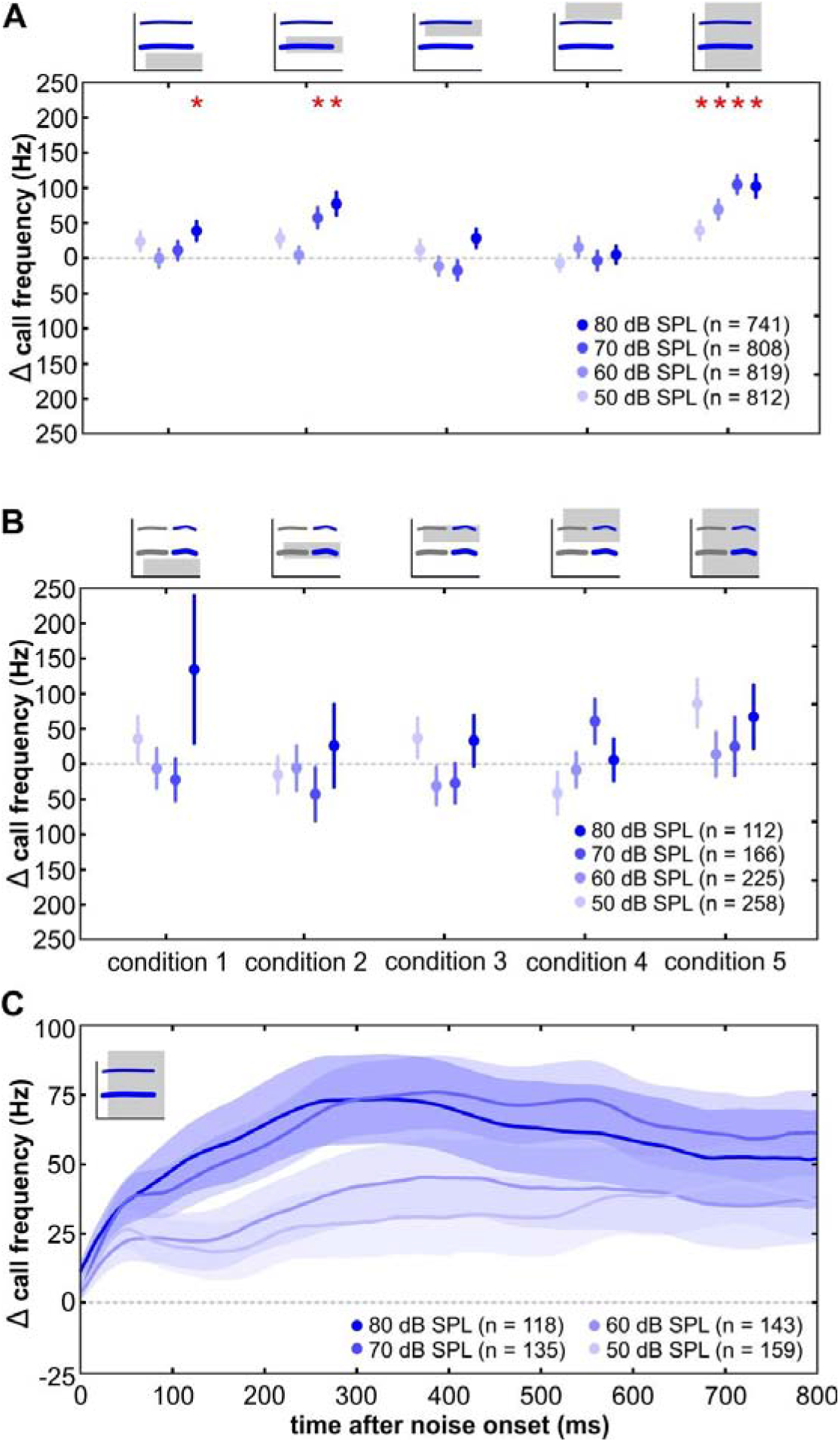
Increasing frequency shifts in response to noise bursts. Δ call frequency (Hz) per corresponding noise condition normalized to control data (dashed lines). Mean of median frequencies after noise onset of each call pooled over all monkeys ± SEM (**A**) for first phee syllables 0–800 ms after noise onset (**B**) for second phee syllables 100–400 ms after second syllable onset. (**C**): First syllables mean Δ call frequency courses (Hz) ± SEM of all amplitude conditions during broadband noise over time after noise onset (ms). Asterisks denote significant differences.

Next, we quantified the magnitude of the observed frequency shifts by calculating population effect sizes (ES) of the factors frequency (ES_freq_), amplitude (ES_ampl_), and the combination of both conditions (ES_freq × ampl_) according to Cohen (1992) (see Material and Methods). An effect would be given if the corresponding ES value of a factor was above the threshold of 0.02 as suggested by Cohen (1992). We found ES_freq × ampl_ values of 0.035 for first syllables and 0.019 for second syllables, indicating an effect for first syllables (**Fig. 2A and 2B**). ES_freq_ for the first syllable was above the threshold (ES_freq_=0.023), while ES_ampl_ was below (ES_ampl_=0.01), indicating that the shifts in fundamental frequency were mainly correlated with the different noise rather than amplitude conditions.

We then tested how fast fundamental frequency shifts occurred within the first phee syllables after noise onset. Therefore, we plotted the mean fundamental frequency courses starting at noise onset times (**Fig. 2C and fig. S5**). The shortest latency of fundamental frequency shifts within a noise condition was defined as the moment where fundamental frequency shifts were significant for a minimum of five consecutive milliseconds after noise onset. Shortest latencies were found for the 0.1–5.1 kHz noise condition at 80 dB (33 ms, n=168) and all broadband conditions (50 dB: 29 ms, n=159; 60 dB: 34 ms, n=143; 70 dB: 25 ms, n=135; 80 dB: 25 ms, n=118), resulting in a mean latency of 29.2±1.9 ms.

Subsequently, we investigated how call amplitudes changed in response to noise perturbation. We calculated mean amplitude shifts after noise onset for first and second phee syllables (**Fig. 3A and fig. 3B**). We found a significant decrease in call amplitude for first phee syllables (F(3,3084)=1.01, p=0.39 for amplitude, F(4,3084)=5.3, p=0.0003 for frequency, n=2019). These shifts were significant for the two middle intensity levels of the 0.1–5.1 kHz noise (60 dB: −1.7±0.5 dB p=3.28e-03, n=103; 70 dB: −2.7±0.5 dB, p=8.17e-04, n=119) as well as for the two middle intensity levels of the broadband noise (60 dB: −2.3±0.6 dB, p=5.15e-03, n=93; 70 dB: −2.0±0.6 dB, p=8.59e-04, n=85). However, we could not find any systematic increase in amplitude shifts or significant amplitude shifts in any of the five noise conditions (n=3093; **Fig. 3A**). Furthermore, the combined effect size (ES_freq × ampl_=0.024) was above 0.02 while the effect size for the frequency (ES_freq_=0.014) and amplitude (ES_ampl_=0.007) factors were below 0.02, indicating that noise perturbation of ongoing first syllables has only a small or no effect on amplitude shifts.

**Figure 3:**
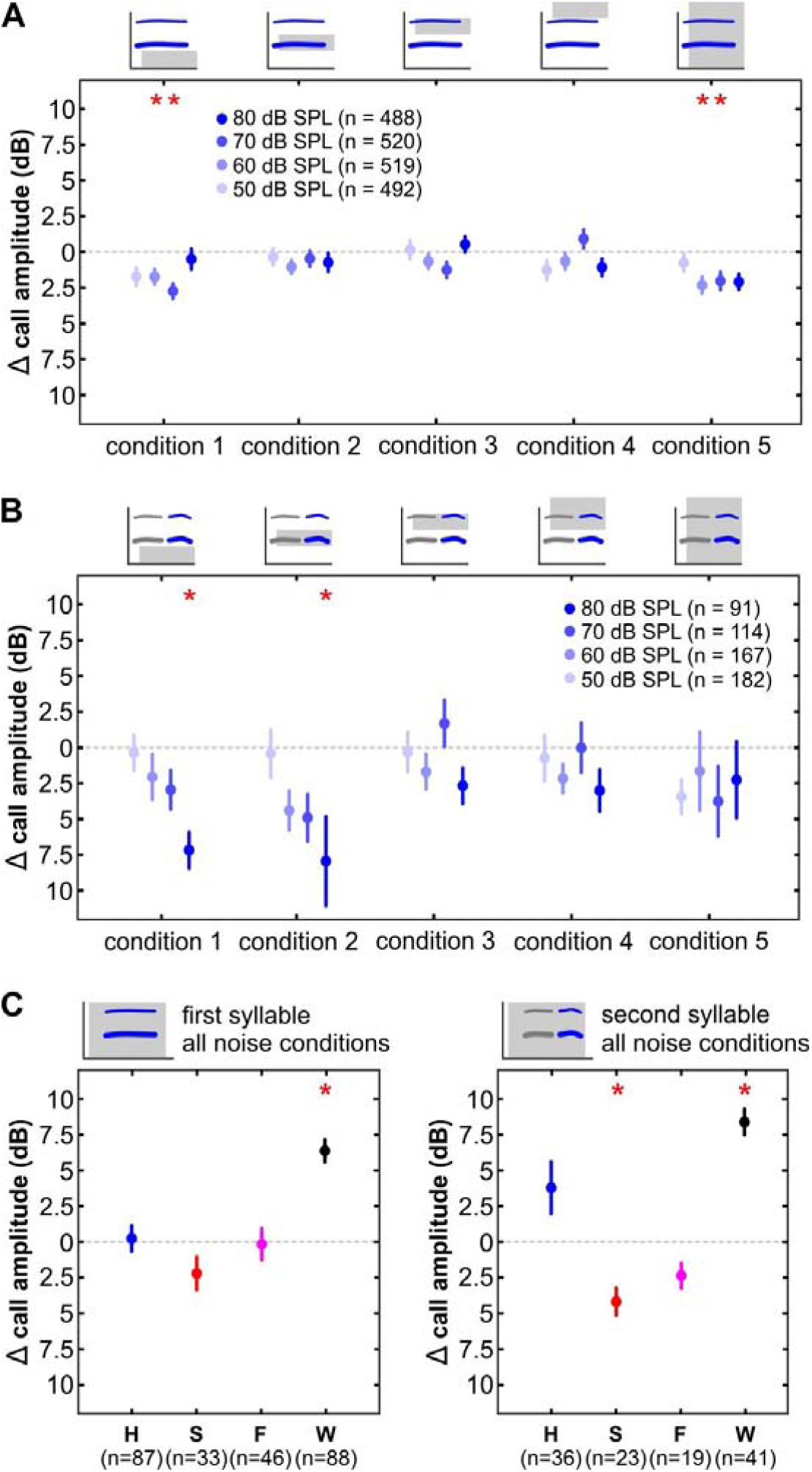
Decreasing amplitude shifts in response to noise bursts. Δ call amplitude (dB) per corresponding noise condition normalized to control data (dashed lines). Mean of median amplitudes after noise onset of each call pooled over all monkeys ± SEM (**A**) for first phee syllables and (**B**) for second phee syllables. Over all noise conditions pooled max Δ amplitudes (dB) ± SEM during 180 s noise per monkey (**C**) for first syllables and (**D**) for second syllables. Asterisks denote significant differences.

However, there was also an amplitude decrease in second phee syllables (F(3,350)=3.76, p=0.011 for amplitude, F(4,950)=1.71, p=0.15 for frequency, n=554). The amplitude shifts in the 0.1–5.1 kHz and 5–10 kHz noise conditions were significant at the highest intensity levels (−7.2±1.3 dB, p=3.90e-02, n=19 and −7.9±3.1 dB, p=2.68e-03, n=16, respectively; **fig. 3B**). Monkeys decreased their call amplitudes in these two conditions with increasing noise intensity levels while no significant call amplitude changes were observed in the other three conditions. All three ES values were above 0.02 (ES_freq × ampl_=0.064, ES_freq_=0.030, ES_ampl_=0.024) suggesting an effect of specific noise perturbation on amplitude shifts of second phee syllables in marmoset monkeys. Although it has been already shown that marmoset monkeys show the Lombard effect while producing twitter calls[17], our results might indicate that marmoset monkeys do not exhibit this reflex when producing phee calls or suppress it and lower their call intensities instead.

To test whether our animals are able to show a Lombard effect or suppress it in a noisy environment in general when producing phee calls, we modified our behavioral experiment scheme. We played back all five noise conditions [0.1–5 kHz, 5–10 kHz, 10–15 kHz, 16–21 kHz, and broadband] at 70 dB SPL amplitude intensity plus two control conditions with a duration of 180 s each, resulting in a block of seven pseudorandomized playback conditions with a total duration of 1260 s. In this new experiment our monkeys produced a total of 803 phee calls (monkey F = 222 phees, H = 270, S = 158, W = 153), which were more commonly uttered (F = 82.5%, H = 80.4%, S = 84.0%, W = 100%) than other produced call types. The relative amounts of single phees ranged between 34.8% and 56.3% and the relative amounts of double phees ranged between 43.71% and 59.49%. Multi-syllabic phees (F = 0.5%, H = 1.9%, S = 5.7%, W = 0%) and segmented phees (F = 0.4%, H = 2.4%, S = 0%, W = 0%) were nearly absent. Monkey H produced 14.3% trill-phees and monkeys F and S produced 15.2% and 13.8% tsik-ekks, respectively. All other call types were below 2.5% for all monkeys. Under these experimental conditions we found that monkey W significantly increased its call intensity for both phee syllables when perturbed by noise (first syllable: 6.4±0.8 dB, p=1.57e-03, n=107; second syllable: 8.4±0.9 dB, p=6.52e-03, n=46; **fig. 3C and fig. S6**), thus, exhibiting the Lombard effect. Furthermore, monkey S significantly decreased the intensity of the second phee syllable and exhibited no changes in call intensity of the first syllable (second syllable: −4.2±1.0 dB, p=2.15e-03, n=52; first syllable: p=0.10, n=68) while monkeys F and H showed no significant amplitude change under noise perturbation (first syllable (H): p=0.89, second syllable (H): p=0.15, n=234; first syllable (F): p=0.91, second syllable (F): p=0.06, n=184). Taken together, the present results suggest that marmosets are capable of exhibiting as well as actively suppressing the Lombard effect in a noisy environment during phee call production.

## Discussion

Our results demonstrate that marmoset monkeys show rapid modulation of call parameters in response to perturbing noise bursts presented after call onset. Ongoing phee vocalizations perturbed by ambient noise rapidly increased call frequency in cases where the fundamental frequency was above or directly masked by the perturbing noise. Bandpass-filtered noise bursts, which did not mask but were above the fundamental frequencies of the calls, had no effect on call frequency. Additionally, call amplitudes of phee calls were affected by low frequency noise bands and broadband noise. Surprisingly, phee calls perturbed after call onset did not exhibit a Lombard effect as previously reported for calls that were produced in constantly presented ambient noise [17,25]. Instead, our monkeys decreased their call intensity in a stepwise function with increasing noise intensity. Our findings suggest a general strategy of avoiding calling in a noisy environment in marmoset monkeys.

### Effects of ambient noise on call frequency

Noise-dependent shifts in call frequency are not well-studied and relatively poorly understood. Only a few studies have reported a rise in call frequencies with increasing amplitudes of ambient noise in birds and bats [8,9,26] and only one study investigated the effect of different noise bands on call frequencies. In bats, the frequencies of echolocation calls increased significantly for a variety of noise stimuli no matter whether they were directly masking the call’s fundamental frequency or presented below the dominant call frequency [8]. In contrast, the present study shows that in marmosets, call frequency was predominantly only affected when we directly masked the calls fundamental frequency. As a result, the strongest rise in call frequencies were found for high noise amplitudes. These findings suggest that the observed rises in call frequencies are an audio-vocal mechanism elicited to increase call detectability in a noisy environment, as has been found in previous studies involving birds [27–29]. Here, it has been predicted that shifts in song frequencies of about 200 Hz increase call detectability by about 10 to 20% [28], which is mainly due to the fact that the spectrum of environmental noise generally shows a decay in amplitude with increasing frequency [28–31]. In the present study, shifts in call frequency occurred with a mean latency of about 30 ms after noise onset suggesting a rapid underlying neural mechanism for frequency modulation. Such fast responses to ambient noise have yet only been found in echolocating bats, which exhibit an increase in call amplitude in about 30 ms after noise onset as well [32].

### Effects of ambient noise on call amplitude

Despite the positive effect of rises in call frequency on signal detectability, the most effective mechanism to improve signal to noise ratio in a noisy environment during vocal production is the Lombard effect, i.e., the involuntary rise in call amplitude as a response to masking noise [12,13]. In the present study, noise perturbation starting after phee call onset had no systematic effect on call amplitude of the first syllable, i.e., the syllable during which noise perturbation started. In cases in which significant shifts occurred, call amplitude did not increase, as expected, but decreased with small effect sizes. This effect was stronger for the second syllables of the phee calls, in which a strong decrease in call amplitude could be observed for low frequency noise conditions. Consequently, call intensity decreased in a stepwise function with increasing noise intensity suggesting a direct effect of noise intensity on call amplitude. In contrast to our study, the Lombard effect has been observed in marmoset monkeys in a previous study [17]. This apparent discrepancy might be explained by the different call types that were investigated in both studies. While we focused on phee calls, a high amplitude call that is produced at the upper end of the amplitude scale [16], the earlier study investigated the twitter call, a vocalization that is produced at lower amplitude intensities [17].

Our results suggest an audio-vocal integration mechanism in marmoset monkeys that is capable of counteracting the Lombard effect. Such a mechanism has been already shown to exist in vocal production learners such as birds and humans [33–36] and seems to be mainly driven by higher-order cognitive processes including cortical structures [13].

### Vocal flexibility during perturbing noise in marmoset monkeys

Current studies have revealed a high degree of vocal flexibility in marmoset monkeys [37], allowing them to control when[14], where[38], and what to vocalize[39]. In addition, recent studies revealed that marmosets are able to modulate distinct call parameters in response to acoustic feedback [18,40]. This vocal flexibility allows marmosets to avoid calling in the presence of environmental noise and predominantly initiate their vocalizations in silent periods [14]. In a previous study, we demonstrated that marmosets interrupt their vocalizations shortly after noise onset when perturbation starts after vocal onset [18], supporting the idea that these animals tend to avoid calling in ambient noise. Such call interruptions, however, were rare (2.6% of all calls), indicating stark neuronal and/or anatomical constraints in exhibiting such behavior [18] and resulting in a large fraction of phee calls being perturbed by noise bursts. In the present study, we show that the call amplitude of such vocalizations are lower.

We suggest that marmoset monkeys exhibit this vocal behavior in a noisy environment to reduce the physiological costs of high intensity phee calls. Marmoset phee calls are elicited at high intensities above 100 dB SPL, resulting in high muscle tensions encompassing almost the entire animal’s body during call production (own observation). Therefore, mechanisms might have evolved in these animals that ensure the proper transmission of these high energetic calls resulting in calling in silent gaps and decreasing call intensity in situations in which sufficient detectability might be potentially diminished, such as during the presence of ambient noise.

### Mechanisms counteracting involuntary audio-vocal effects need cognitive control

Based on the current work and earlier studies [14,18], we propose a hypothetical neuronal model suggesting various audio-vocal control mechanism involving cortical, subcortical, and corticofugal connections capable of modulating vocal behavior in a noisy environment (**Fig. 4**). In accordance to earlier work [41,42], our model consists of a volitional articulatory motor network originating in the prefrontal cortex (PFC) cognitively controlling vocal output of a phylogenetically conserved primary vocal motor network predominantly consisting of a subcortical neuronal network. The vocal motor network can be modulated by auditory structures on several cortical and subcortical brain levels [13]. The decision to initiate or suppress a call, as well as counteracting an involuntary effect (Lombard effect), needs cognitive control. The ability to interrupt calls or modulate call parameters as a response to perturbing noise might be controlled by both subcortical mechanisms and corticofugal projections. Neurophysiological studies will now have to elucidate at which brain levels audio-vocal integration mechanisms exist that explain the observed capabilities of marmoset monkeys to counteract a previously thought involuntary audio-vocal mechanism, the Lombard effect.

**Figure 4:**
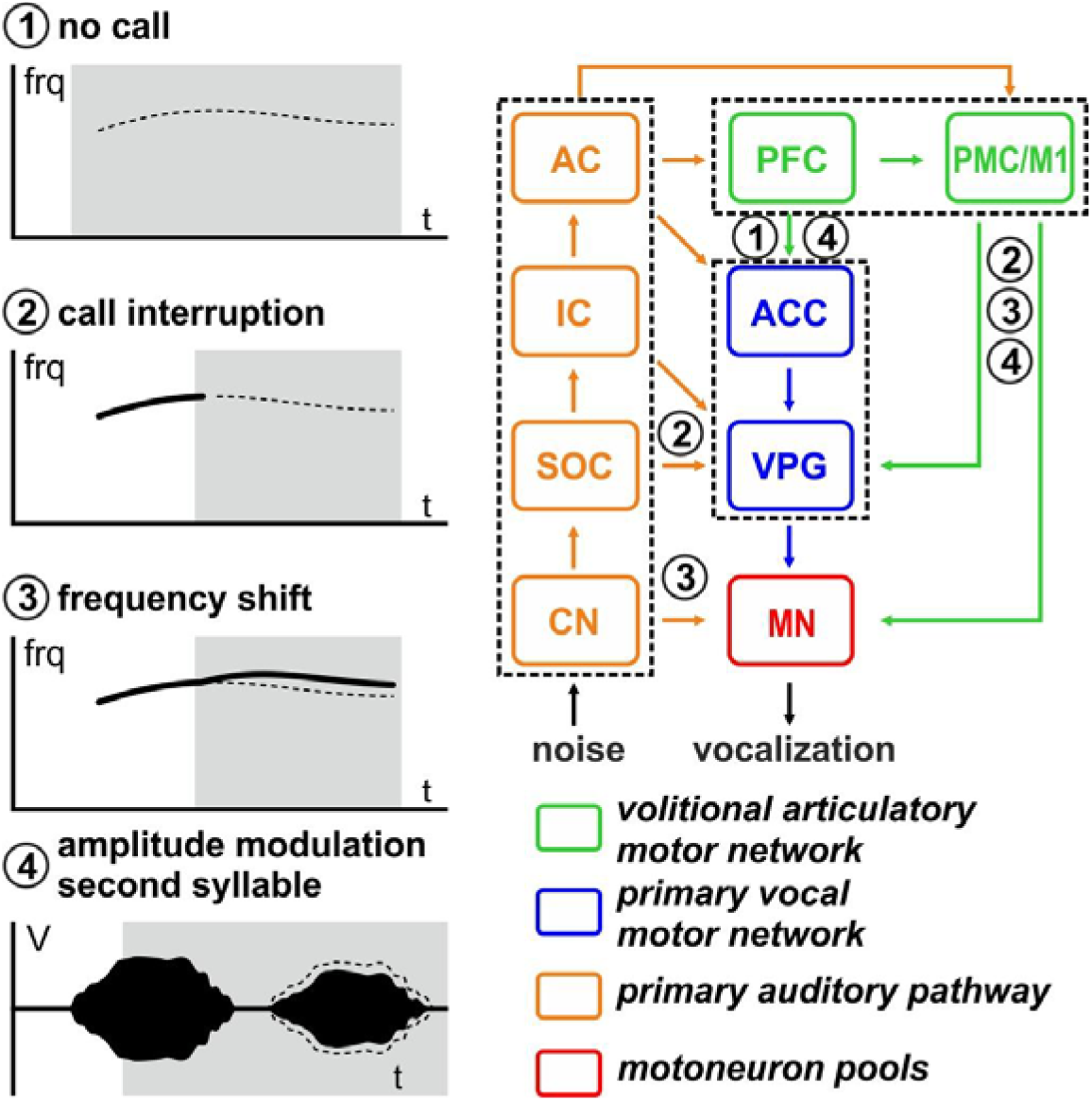
Hypothetical neuronal model for audio-vocal interaction. Call production might be affected by ambient noise at different brain levels. Audio-vocal integration mechanisms are known to happen between cortical and subcortical structures as well as via corticofugal projections. See text for further explanation.

## Material and Methods

### Animal Housing and Maintenance

Four adult marmoset monkeys (*Callithrix jacchus*) were used in the present study. Monkeys were usually kept in different sex pairs and were all born in captivity. The animals had *ad libitum* access to water and were fed on a restricted food protocol including a daily basis of commercial pellets, fruits, vegetables, mealworms, and locusts. Additional treats, such as marshmallows or grapes, were used as positive reinforcements to transfer the animals from their home cage to the experimental cage. Environmental conditions in the animal husbandry were maintained at a temperature of 26°C, 40-60% relative humidity, and a 12h:12h day/night cycle. All animal handling procedures were in accordance with the guidelines for animal experimentation and authorized by the national authority, the Regierungspräsidium Tübingen. All vocalizations analyzed in this study are a fraction of calls that have been recorded in a previous study (Pomberger et al. 2018).

### Experimental Setup and Procedure

The vocal behavior of four animals was recorded in a soundproof chamber in response to noise playback that was initiated after vocal onset as reported earlier (Pomberger et al., 2018). Briefly, the animals were transferred into a recording cage (0.6 × 0.6 × 0.8 m), which was placed in a soundproofed chamber, with *ad libitum* access to water and food pellets throughout the recording period. In this behavioral setup, marmoset monkeys predominantly produce phee calls to interact with conspecifics (phee ratio within all uttered calls; monkey S: 99.1 %, H: 92.0 %, W: 95.6 %, F: 96.8 %). Other call types such as trill-phees, twitter, trills, tsik-ekks [22], or segmented phees [24] were only rarely uttered (ratios were well below 2.5% for all other call types in all monkeys except trill-phees in monkey H [4.6 %]). Monkeys produced a mean of 118±9 (monkey S), 167±31 (H), 117±10 (W), and 87±7 (F) phee calls per session. The vocal behavior of each individual monkey was recorded once a day in sessions ranging between one and two hours in duration. Data were collected in sessions at various times during the day between 11 am and 5 pm. Recordings were performed for 10–28 days (mean: 17±3 days) for each individual animal. The monkey’s behavior was constantly monitored and observed with a video camera (ace acA1300-60gc, Basler, Germany with 4.5–12.5 mm CS-Mount Objective H3Z4512CS-IR 1/2, Computar, Japan) placed on top of the cage and recorded with standard software (Ethovision XT version 4.2.22, Noldus, the Netherlands). The vocal behavior was collected with eight microphones (MKH 8020 microphone with MZX 8000 preamplifier, Sennheiser, Germany), which were positioned in an octagonal design around the cage (**Fig. 1A**), digitized using an A/D interface (Octacapture, Roland, Japan; sample rate: 96 kHz), and recorded using standard software (Avisoft-Recorder, Avisoft Bioacoustics, Germany). A custom-written program (OpenEX, Tucker-Davis Technologies, U.S.A.) running on a workstation (WS-X in combination with an RZ6D multi I/O processor, Tucker-Davis Technologies, U.S.A.) monitored the vocal behavior in real-time via an additional microphone (MKH 8020 microphone with MZX 8000 preamplifier, Sennheiser, Germany) placed on top of the cage, which automatically detected vocalizations through online calculation of several acoustic parameters, such as call intensity, minimum duration of call intensity duration, call frequency, and several spectral features. The median vocal detection rate was well above 99% and three out of four vocalizations were detected within the first 146 ms after call onset (**Fig. 1B**).

The eight microphones positioned around the cage were installed to ensure precise calculation of dB SPL values of vocalizations with a corresponding microphone being positioned in front of the monkey (for detail see below).

For two out of three uttered vocalizations, we played back noise bursts of different frequency-bands and amplitudes via a loudspeaker (MF1 Multi-Field Magnetic Speakers, Tucker-Davis Technologies, U.S.A.) positioned on top of the cage, immediately after vocal detection. Noise bursts had a duration of 4 s (including 10 ms rise times) to ensure noise perturbation throughout the first phee syllable as well as the initiation of the second syllable (**Fig. 1C**). Five different noise band conditions (broadband noise and bandpass filtered noise bands: 0.1–5.1 kHz, 5–10 kHz, 10–15 kHz, and 16–21 kHz) were played back at four different amplitudes (50 dB, 60 dB, 70 dB, 80 dB) each. All 20 noise conditions were played back pseudo-randomly in blocks of 30 uttered vocalizations, resulting in 20 calls being perturbed with noise after call onset and 10 calls without noise playback remaining unaffected (control). After one block ended, a new block was generated. Noise playback generation and presentation were performed with the same custom-written software used for call detection.

### Data Analysis

We programmed a custom-written GUI (Matlab, Mathworks, U.S.A.) to clock Avisoft, Noldus, and TDT recordings offline and to extract the detected calls from the recording channel with the best SNR. Vocal onset to offset were manually flagged as well as noise onset times using the aligned sono- and spectrogram of vocalizations. We used a Hanning window with a 512-window size, 1024 FFT, overlap of 25%, and temporal resolution of one millisecond. We only considered first phee syllables for calculation that were detected/perturbed within 200 ms of call onset and with a minimum duration of 800 ms. Consequently, first phee syllables that were interrupted directly after noise onset as previously reported in an earlier study [18] were excluded from further analysis. Second phee syllables were only analyzed if they had a minimum duration of 500 ms. In rare cases, call termination could not be visually detected due to overlapping noise (mostly during the 80 dB SPL condition). These calls were also excluded from further analysis.

After labelling a call, peak frequencies of the fundamental component were automatically calculated within one-millisecond time bins (8192 FFT, 96 kHz sample rate resulting in a frequency resolution of 11.71 Hz). In cases where the SNR between the call amplitude and playback noise was not high enough for automatic fundamental peak frequency calculation, frequency trajectories were calculated by manually setting call frequencies at several time points and interpolating call frequencies in between the set values. The accuracy of manual labelling compared to automatic calculation of peak frequencies was high and median differences between both techniques below the frequency resolution used (**Fig. S1)**.

Call amplitudes were calculated for all phee calls during which the animals did not move their heads during call production. For these calls, head positions were manually labelled by marking the two white ear tufts in the GUI (see **Fig. 1B**). Next, a perpendicular line starting at the center of the later connection was used to compute angles of the microphones indicating the monkey’s relative head position. The microphone with the smallest angle to the perpendicular line was used for further calculation (**Fig. 1B**). Calls that were uttered in the rare cases where the angle between the front of the monkey’s head and the microphone was more than 45 degrees were excluded from further analysis. Furthermore, phee calls that were uttered during head movements of the animal were not used for amplitude calculations and only considered for fundamental frequency calculation resulting in a larger data set for frequency analysis.

From the recordings of the microphone foremost in front of the animal, call amplitude trajectories (in dB SPL) were calculated using a sliding window approach (window size: 25 ms; step length: 1 ms). Sound levels of the recorded playback noise were determined for all conditions and subtracted from the call amplitude measurements taken, using a modification of the spectral noise subtraction method [43]. Briefly, we first calculated an estimate for each noise band by calculating the mean of ten recordings of one noise condition for each microphone. Then, we subtracted this noise estimate in the spectral dimension from noise perturbed parts of a call (i.e., from noise onset to the end of the call) and corrected the outcome as shown in formula (1), where *P*_*s*_(*w*) is the spectrogram of the signal and the noise, *P*_*n*_(*w*) the spectrogram of the noise estimate and *P*_*s*_′(*w*) the modified signal spectrum. Alpha is defined as the subtraction factor and beta as the spectral floor parameter.

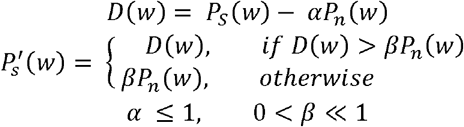

Alpha and beta were calculated using the following equation:

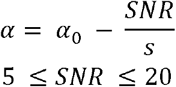

According to Berouti et al. [43], we chose alpha_0_ = 4 and s = 20/3 as a best fit for proper amplitude calculation. A simple empirical test verified the method; a control phee was played and recorded in the recording chamber ten times with broadband noise 70 dB SPL, ten times with a 5–10 kHz noise band and ten times under control conditions (no noise). As reported previously, differences between conditions of <1 dB can be assumed to be negligible [44]. In our case, median differences between control and both noise conditions were below 1 dB (broadband: 0.8 dB, 5–10 kHz noise band: 0.3 dB; **Figs. S2A and B**) indicating successful performance of the used method. The distance of the animal’s head to the microphone was considered by adding a distance factor directly after noise subtraction to the measurements resulting in a standardized amplitude trajectory (in dB) of each call as produced 10 cm in front of the animal’s head.

### Frequency/amplitude calculation and normalization

Mean fundamental frequency values were obtained with a sliding window approach (window size: 10 ms, step size: 1 ms). We then calculated the mean of the fundamental frequency in a 20-5 ms time window prior to noise onset (for the noise conditions) and call detection (for the control condition) for each individual call and subtracted this value from each of the frequency values after noise onset. Finally, all values of calls in the noise conditions were normalized by subtracting the mean of the respective frequency value of the control condition. Amplitude values were calculated in a similar way. Here, we also calculated the mean amplitude for each individual call in a 20-5 ms time window prior to noise onset and subtracted these values from the mean amplitude values after noise onset. According to the frequency normalization, we then normalized all amplitude values by subtracting the mean of the amplitude values from the corresponding values in the control condition. For the 180 s noise experiment, we used the calculated amplitude values as described above in *data analysis*.

### Phee call discrimination models

Marmoset monkeys tend to interrupt their phee calls after the first syllable in response to noise perturbation [18]. For perturbed phee calls, we consequently assumed that a substantial number of single phees had to be interrupted double phees. Recently, it has been shown that single phees and the first syllables of double phees significantly differ in a number of call parameters, such as call frequency and duration [20]. We therefore had to find a way to distinguish single phee calls that were interrupted double phees from original single phees prior to data normalization. To address this, we used the findings of Miller and colleagues [20] that suggested that early peak frequencies and durations of phee calls are sufficient to predict whether a phee call consists out of one or two syllables. Additionally, we found that this is also true for early amplitude values of a call. We applied a quadratic classification model (MATLAB) to discriminate between single and double phees with a two-dimensional classifier for fundamental frequency analysis using 1st syllable durations and peak frequencies at 25 ms after call onset for frequency analyses (**Fig. S3**). Since we observed that early amplitude values are also a good predictor (**Fig. S4**), we used a three-dimensional classifier with call amplitude values at 25 ms after call onset as the third measure for amplitude analyses (**Fig. S4**). Basically, in a first step the mean, μ_k_, and covariance matrix, Σ_k_, of each class is calculated from all control values to obtain the density function of the multivariate normal at a point, x, using the following formula:

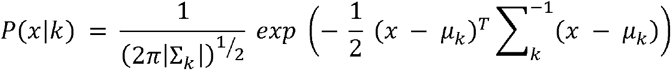

Where |Σ_k_| is the determinant of Σ_k_, and 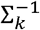 is the inverse matrix. Using the prior probability *P*(*k*) of class *k* and *P*(*x*) as a normalization constant we obtain the posterior probability 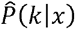 that a point *x* belongs to class *k* based on:

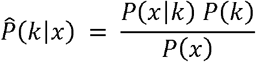

These results are then used to classify our phee calls into single and double phees by minimizing the expected classification cost using:

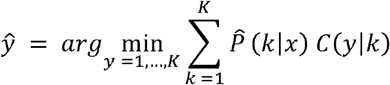

Where 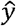 is the predicted classification, *K* is the number of classes,and *C*(*y*|*k*) is the cost of classifying an observation as y when its true class is k. In total, the loss for the 2D classification was between 10.8% and 23.2% (mean: 15.1±2.8) and for the 3D classification between 6.8% and 15.7% (mean: 12.3±1.9) for each monkey.

### Statistical analysis

Statistical analyses were performed with MATLAB (2016b, MathWorks, Natick, MA). We performed a two-way ANOVA to test for significant differences in shifts of fundamental call frequency and amplitude within all noise band conditions (alpha = 0.05, Bonferroni corrected). Effect sizes (ES) were calculated using the following formula:

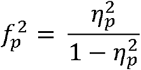

Where 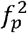 represents the effect size of factor ***p*** and 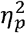 is calculated as:

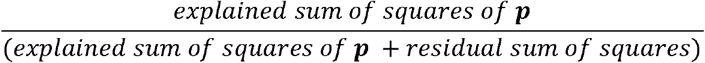

### Data availability

All data needed to evaluate the conclusions in the paper are present in the paper. Additional data related to this paper may be requested from the corresponding author.

## Author Contributions

S.R.H. conceived the study; T.P. and S.R.H designed the experiments; T.P. and J.L. conducted the experiments and performed data analyses; all authors interpreted the data and wrote the manuscript. S.R.H. provided the animals and supervised the project.

## Acknowledgments

We thank John Holmes for proofreading. This work was supported by the Werner Reichardt Centre for Integrative Neuroscience (CIN) at the Eberhard Karls University of Tübingen (CIN is an Excellence Cluster funded by the Deutsche Forschungsgemeinschaft within the framework of the Excellence Initiative EXC 307).

## Supplemental Figures

**Figure S1:**
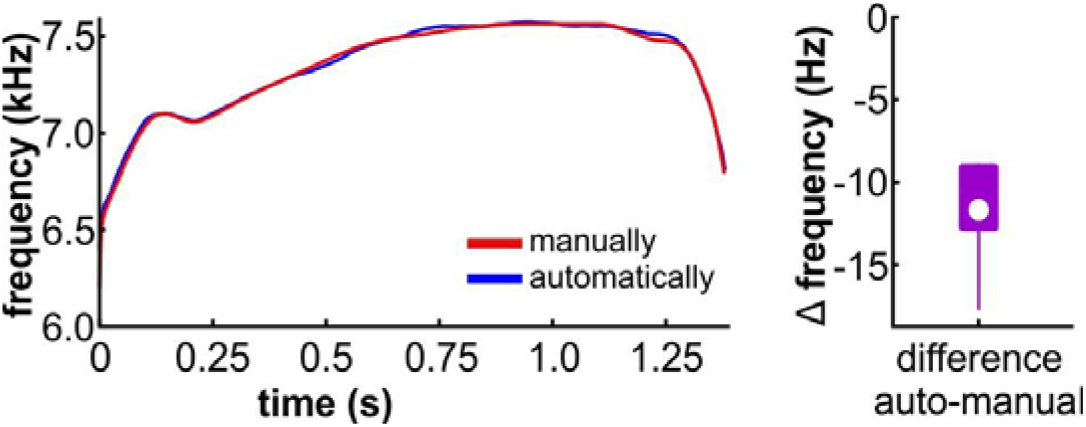
Comparison of automatically vs manually marked calls. (**A**) Frequency course (Hz) over time (ms) of a manually (blue) and automatically (red) marked example phee call. (**B**) Mean Δ frequencies of automatically-manually marked calls.

**Figure S2:**
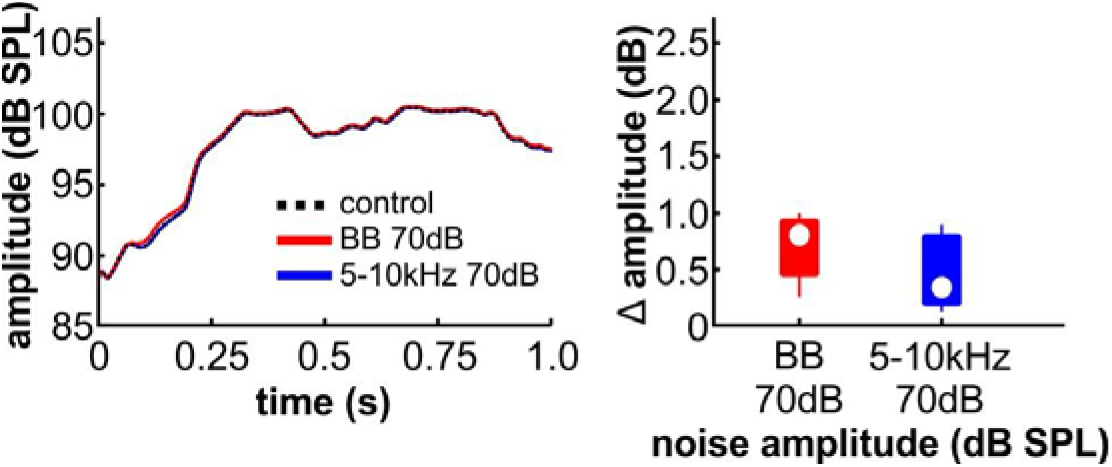
Test for noise subtraction accuracy. (**A**) Mean amplitude courses (dB SPL) for both 70 dB overlapping noise conditions (5–10 kHz, blue; broadband, red) as well as for the control (no noise, black dashed) of 10 test phees over time (ms). (**B**) Maximum Δ amplitude (dB) compared to control calls of 70 dB broadband and 5–10 kHz noise conditions.

**Figure S3:**
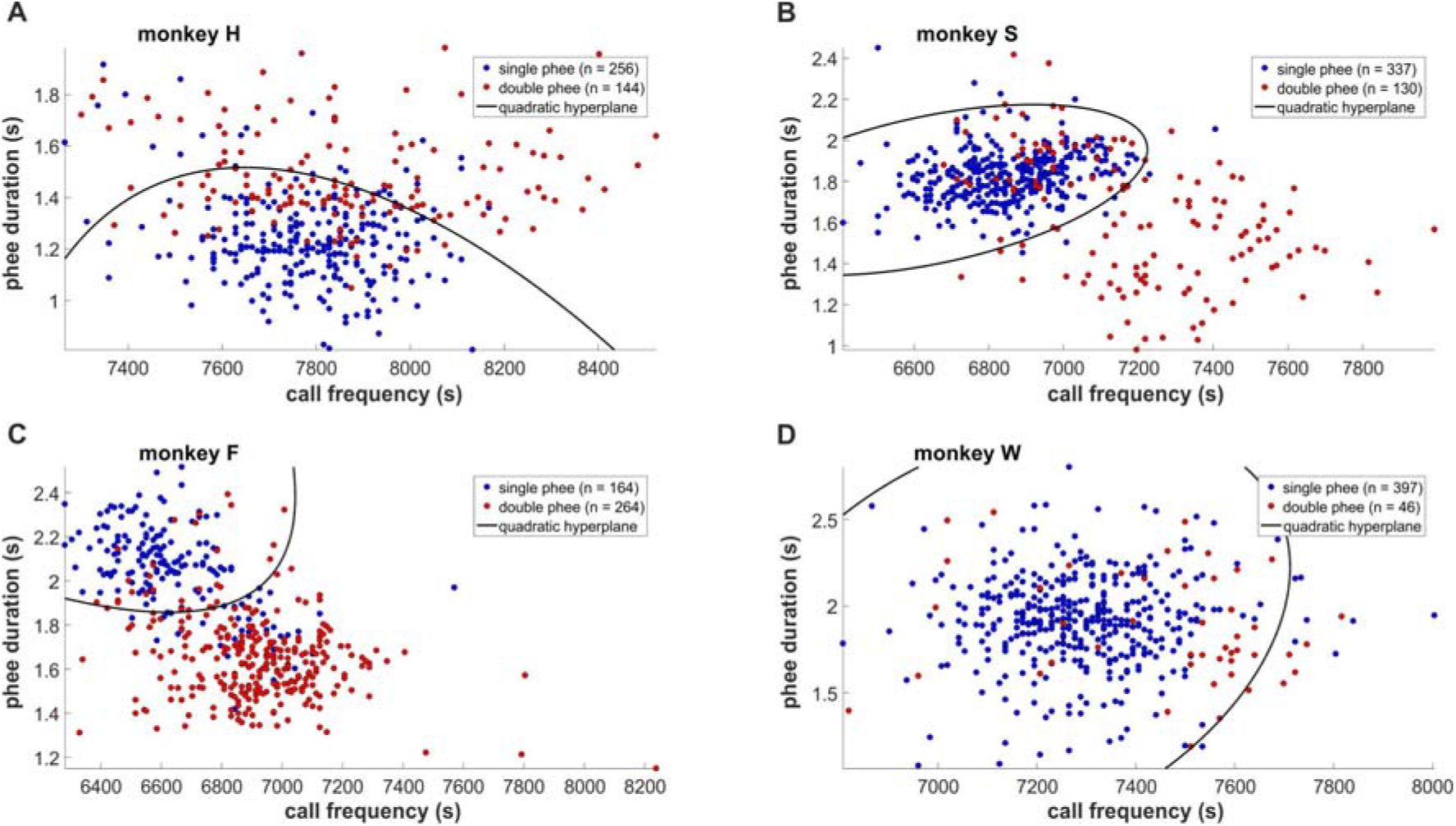
Single-double phee discrimination with a two-dimensional classifier. For single-double phee discrimination, first syllable fundamental peak frequencies (Hz) at 25 ms after call onset and corresponding phee durations (s) were used.

**Figure S4:**
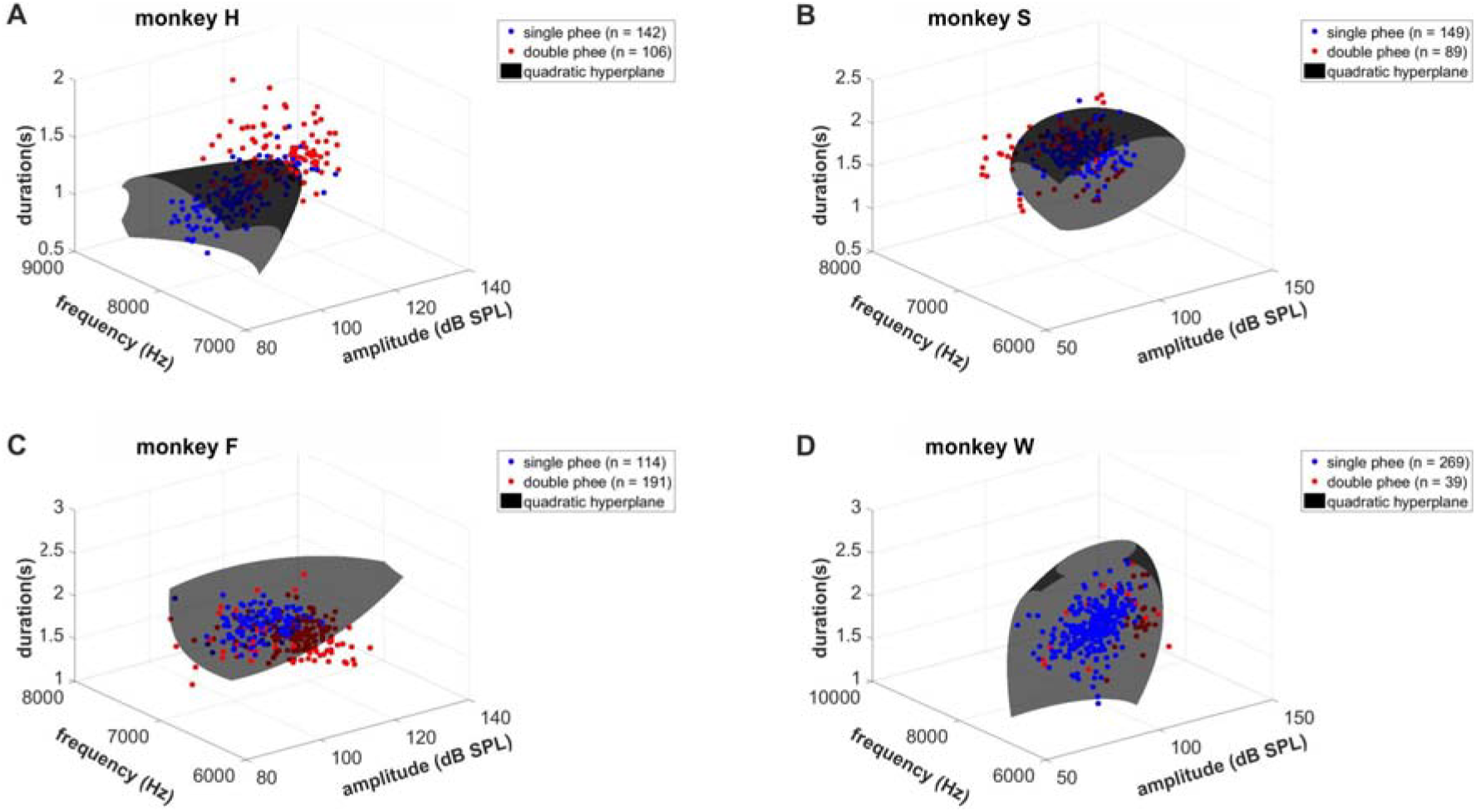
Single-double phee discrimination with a three-dimensional classifier. For single-double phee discrimination, first syllable fundamental peak frequencies (Hz), as well as amplitudes (dB SPL) at 25 ms after call onset and corresponding phee durations (s), were used.

**Figure S5:**
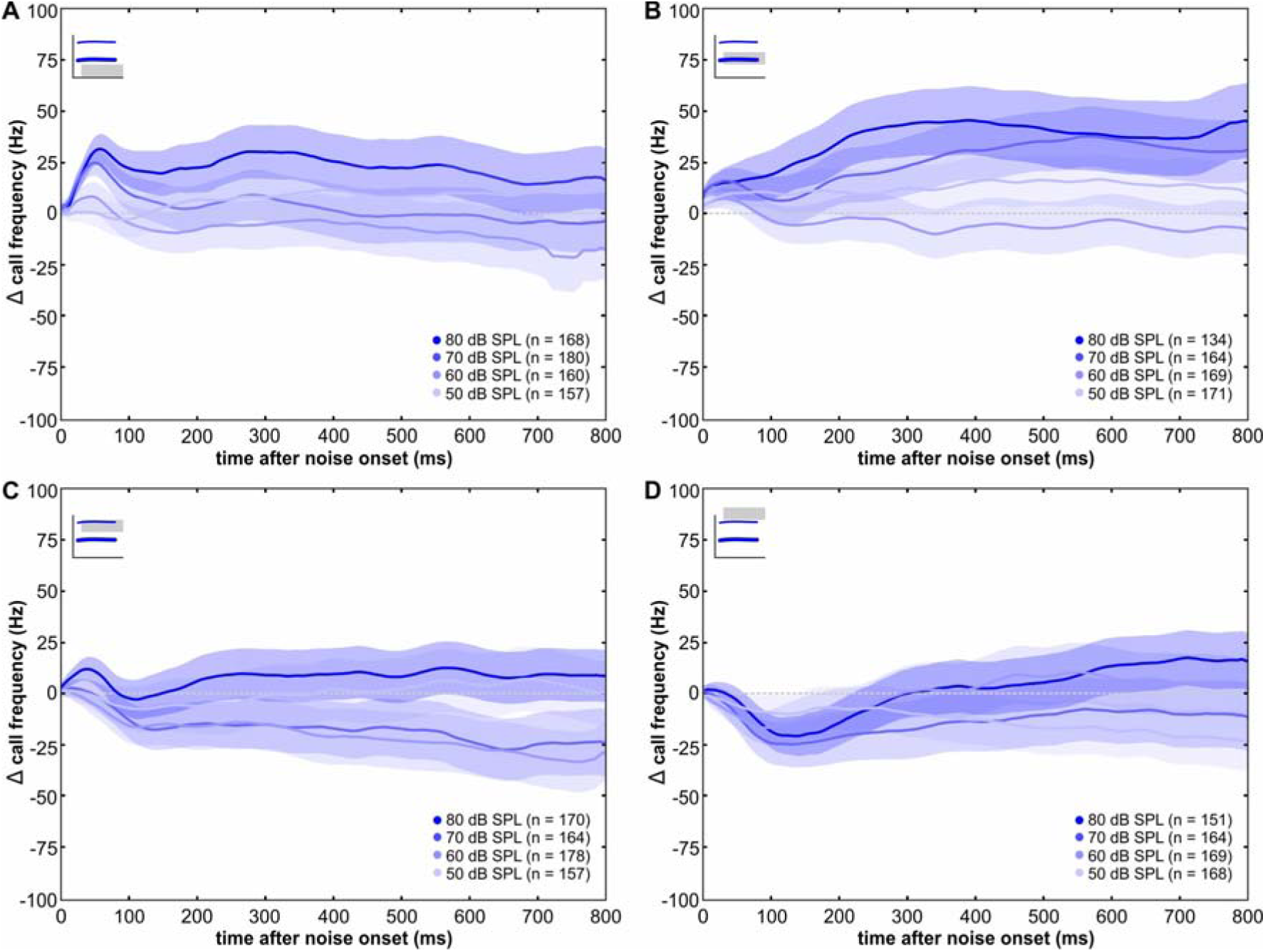
Frequency courses in response to noise bursts. First syllables mean Δ frequency courses (Hz) ± SEM of all amplitude conditions normalized to control data (dashed lines) during (**A**) 0.1–5.1 kHz, (**B**) 5–10 kHz, (**C**) 10–15 kHz, and (**D**) 16–21 kHz noise conditions over time after noise onset (ms).

**Figure S6:**
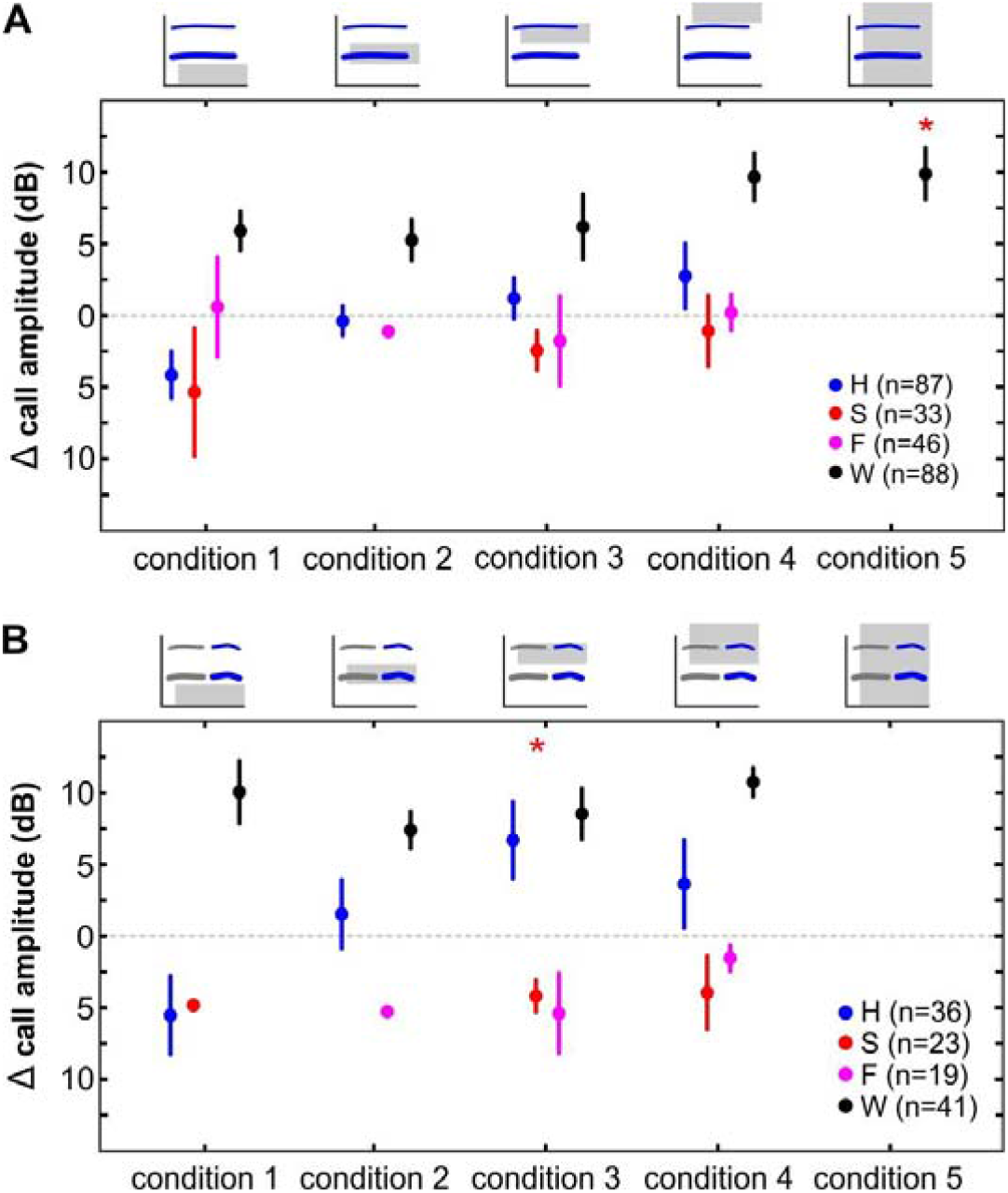
Amplitude shifts in response to noise bursts. Pooled maximum Δ amplitudes (dB) ± SEM during 180 s noise per monkey per noise condition normalized to control data (dashed lines) (**A**) for first syllables and (**B**) for second syllables. Asterisks denote significant differences.

## References

1. Poldrack RA, Farah MJ. 2015 Progress and challenges in probing the human brain. Nature 526, 371–379.

2. Stockhorst U, Pietrowsky R. 2004 Olfactory perception, communication, and the nose-to-brain pathway. Physiol. Behav. 83, 3–11.

3. Osorio D, Vorobyev M. 2008 A review of the evolution of animal colour vision and visual communication signals. Vision Res. 48, 2042–2051

4. Ackermann H, Hage SR, Ziegler W. 2014 Brain mechanisms of acoustic communication in humans and nonhuman primates: An evolutionary perspective. Behav. Brain Sci. 37, 529–546.

5. Bradbury JW, Vehrencamp SL. 1998 Principles of Animal Communication.

6. Brumm H, Slabbekoorn H. 2005 Acoustic Communication in Noise. Adv. Study Behav. 35, 151–209.

7. Lombard E. 1911 Le signe de l’elevation de la voix. Ann. Mal. L’Oreille du Larynx XXXVII, 101–119.

8. Hage SR, Jiang T, Berquist SW, Feng J, Metzner W. 2013 Ambient noise induces independent shifts in call frequency and amplitude within the Lombard effect in echolocating bats. Proc. Natl. Acad. Sci. U. S. A. 110, 4063–4068.

9. Osmanski MS, Dooling RJ. 2009 The effect of altered auditory feedback on control of vocal production in budgerigars (*Melopsittacus undulatus*). J. Acoust. Soc. Am. 126, 911–919.

10. Brumm H. 2006 Signalling through acoustic windows: Nightingales avoid interspecific competition by short-term adjustment of song timing. J. Comp. Physiol. A Neuroethol. Sensory, Neural, Behav. Physiol. 192, 1279–1285.

11. Luo J, Goerlitz HR, Brumm H, Wiegrebe L. 2015 Linking the sender to the receiver: Vocal adjustments by bats to maintain signal detection in noise. Sci. Rep. 5, 1–11.

12. Brumm H, Zollinger A. 2011 The evolution of the Lombard effect: 100 years of psychoacoustic research. Behaviour 148, 1173–1198.

13. Luo J, Hage SR, Moss CF. 2018 The Lombard Effect: From Acoustics to Neural Mechanisms. Trends Neurosci. 41, 938–949.

14. Roy S, Miller CT, Gottsch D, Wang X. 2011 Vocal control by the common marmoset in the presence of interfering noise. J. Exp. Biol. 214, 3619–3629.

15. Zelick RD, Narins PM. 1982 Analysis of acoustically evoked call suppression behaviour in a neotropical treefrog. Anim. Behav. 30, 728–733.

16. Eliades SJ, Wang X. 2012 Neural correlates of the lombard effect in primate auditory cortex. J. Neurosci. 32, 10737–48.

17. Brumm H. 2004 Acoustic communication in noise: regulation of call characteristics in a New World monkey. J. Exp. Biol. 207, 443–448.

18. Pomberger T, Risueno-Segovia C, Löschner J, Hage SR. 2018 Precise Motor Control Enables Rapid Flexibility in Vocal Behavior of Marmoset Monkeys. Curr. Biol. 28, 788–794.e3.

19. Miller CT, Flusberg S, Hauser MD. 2003 Interruptibility of long call production in tamarins: implications for vocal control. J. Exp. Biol. 206, 2629–2639.

20. Miller CT, Eliades SJ, Wang X. 2009 Motor planning for vocal production in common marmosets. Anim. Behav. 78, 1195–1203.

21. Egnor SER, Iguina CG, Hauser MD. 2006 Perturbation of auditory feedback causes systematic perturbation in vocal structure in adult cotton-top tamarins. J. Exp. Biol. 209, 3652–3663.

22. Agamaite JA, Chang C-J, Osmanski MS, Wang X. 2015 A quantitative acoustic analysis of the vocal repertoire of the common marmoset (Callithrix jacchus). J. Acoust. Soc. Am. 138, 2906–2928.

23. Pistorio AL, Vintch B, Wang X. 2006 Acoustic analysis of vocal development in a New World primate, the common marmoset (*Callithrix jacchus*). J. Acoust. Soc. Am. 120, 1655–1670.

24. Zürcher Y, Burkart JM. 2017 Evidence for Dialects in Three Captive Populations of Common Marmosets (Callithrix jacchus). Int. J. Primatol. 38, 780–793.

25. Egnor SR, Hauser M. 2006 Noise-induced vocal modulation in cotton-top tamarins (Saguinus oedipus). Am. J. Primatol. 68, 1183–1190.

26. Schuster S, Zollinger SA, Lesku JA, Brumm H. 2012 On the evolution of noise-dependent vocal plasticity in birds. Biol. Lett. 8, 913–916.

27. Bermúdez-Cuamatzin E, Ríos-Chelén AA, Gil D, Garcia CM. 2011 Experimental evidence for real-time song frequency shift in response to urban noise in a passerine bird. Biol. Lett. 7, 36–38.

28. Nemeth E, Brumm H. 2010 Birds and Anthropogenic Noise: Are Urban Songs Adaptive? Am. Nat. 176, 465–475.

29. Pohl NU, Leadbeater E, Slabbekoorn H, Klump GM, Langemann U. 2012 Great tits in urban noise benefit from high frequencies in song detection and discrimination. Anim. Behav. 83, 711–721.

30. Halfwerk W, Slabbekoorn H. 2009 A behavioural mechanism explaining noise-dependent frequency use in urban birdsong. Anim. Behav. 78, 1301–1307.

31. Pohl NU, Slabbekoorn H, Klump GM, Langemann U. 2009 Effects of signal features and environmental noise on signal detection in the great tit, Parus major. Anim. Behav. 78, 1293–1300.

32. Luo J, Kothari NB, Moss CF. 2017 Sensorimotor integration on a rapid time scale. Proc. Natl. Acad. Sci. 114, 6605–6610.

33. Kobayasi KI, Okanoya K. 2003 Context-dependent song amplitude control in Bengalese finches. Neuroreport 14, 521–524.

34. Pick HL, Siegel GM, Fox PW, Garber SR, Kearney JK. 1989 Inhibiting the Lombard effect. J. Acoust. Soc. Am. 85, 894–900.

35. Therrien AS, Lyons J, Balasubramaniam R. 2012 Sensory Attenuation of Self-Produced Feedback: The Lombard Effect Revisited. PLoS One 7, e49370.

36. Vinney LA, van Mersbergen M, Connor NP, Turkstra LS. 2016 Vocal Control: Is It Susceptible to the Negative Effects of Self-Regulatory Depletion? J. Voice 30, 638.e21–638.e31.

37. Ghazanfar AA, Liao DA, Takahashi DY. 2019 Volition and learning in primate vocal behaviour. Anim. Behav., 1–9.

38. Choi JY, Takahashi DY, Ghazanfar AA. 2015 Cooperative vocal control in marmoset monkeys via vocal feedback. J. Neurophysiol. 114, 274–283.

39. Liao DA, Zhang YS, Cai LX, Ghazanfar AA. 2018 Internal states and extrinsic factors both determine monkey vocal production. Proc. Natl. Acad. Sci. 115, 201722426.

40. Eliades SJ, Tsunada J. 2018 Auditory cortical activity drives feedback-dependent vocal control in marmosets. Nat. Commun. 9, 2540.

41. Hage SR, Nieder A. 2016 Dual Neural Network Model for the Evolution of Speech and Language. Trends Neurosci. 39, 813–829.

42. Hage SR. 2019 Precise vocal timing needs cortical control. Science. 363, 926–928.

43. Berouti M, Schwartz R, Makhoul J. 1978 Enhancement of speech corrupted by acoustic noise. In ICASSP ’79. IEEE International Conference on Acoustics, Speech, and Signal Processing, pp. 208–211.

44. Brumm H, Schmidt R, Schrader L. 2009 Noise-dependent vocal plasticity in domestic fowl. Anim. Behav. 78, 741–746.

